# Machine learning illuminates how diet influences the evolution of yeast galactose metabolism

**DOI:** 10.1101/2023.07.20.549758

**Authors:** Marie-Claire Harrison, Emily J. Ubbelohde, Abigail L. LaBella, Dana A. Opulente, John F. Wolters, Xiaofan Zhou, Xing-Xing Shen, Marizeth Groenewald, Chris Todd Hittinger, Antonis Rokas

## Abstract

How genomic differences contribute to phenotypic differences across species is a major question in biology. The recently characterized genomes, isolation environments, and qualitative patterns of growth on 122 sources and conditions of 1,154 strains from 1,049 fungal species (nearly all known) in the subphylum Saccharomycotina provide a powerful, yet complex, dataset for addressing this question. In recent years, machine learning has been successfully used in diverse analyses of biological big data. Using a random forest classification algorithm trained on these genomic, metabolic, and/or environmental data, we predicted growth on several carbon sources and conditions with high accuracy from presence/absence patterns of genes and of growth in other conditions. Known structural genes involved in assimilation of these sources were important features contributing to prediction accuracy, whereas isolation environmental data were poor predictors. By further examining growth on galactose, we found that it can be predicted with high accuracy from either genomic (92.6%) or growth data in 120 other conditions (83.3%) but not from isolation environment data (65.7%). When we combined genomic and growth data, we noted that prediction accuracy was even higher (93.4%) and that, after the *<u>GAL</u>*actose utilization genes, the most important feature for predicting growth on galactose was growth on galactitol. These data raised the hypothesis that several species in two orders, Serinales and Pichiales (containing *Candida auris* and the genus *Ogataea*, respectively), have an alternative galactose utilization pathway because they lack the *GAL* genes. Growth and biochemical assays of several of these species confirmed that they utilize galactose through an oxidoreductive D-galactose pathway, rather than the canonical *GAL* pathway. We conclude that machine learning is a powerful tool for investigating the evolution of the yeast genotype-phenotype map and that it can help uncover novel biology, even in well-studied traits.

## Introduction

Yeasts in the subphylum Saccharomycotina (hereafter referred to as yeasts) are genomically diverse, geographically widely distributed, found in diverse habitats, and utilized for diverse purposes by humans – the baker’s yeast *Saccharomyces cerevisiae* is the cornerstone of the winemaking, brewing, baking, and biotech industries; *Candida albicans* is a human commensal that thrives in the human gut and occasionally becomes a serious pathogen; *Candida auris* is an emerging fungal pathogen of great concern because of its innate resistance to available antifungal drugs; and *Lipomyces starkeyi* produces lipids and has several biotechnology applications (Hittinger et al. 2018 Yaguchi et al. 2017, Case et al. 2022).

Yeast ecological diversity is thought to be intimately tied to the vast diversity in their diets, i.e., the diversity of primary metabolic capabilities that allow them to grow on many different sources of carbon and nitrogen (Opulente et al. 2018). However, we currently lack a comprehensive understanding of how variation in yeast gene content or regulation is related to the metabolic diversity and environmental adaptation of the ∼1,200 species found across the subphylum. Recently, the Y1000+ Project (http://y1000plus.org/) published draft genome sequences of 1,086 representative strains (mostly taxonomic type strains) from nearly 1,016 and 62 novel candidate species of yeasts (Shen et al. 2018, Opulente et al. 2023, Hittinger et al. 2015). The Y1000+ Project has also systematically recorded (from the literature) and / or experimentally generated the isolation environments and qualitative and quantitative patterns of growth on diverse carbon sources, nitrogen sources, and environmental conditions (e.g., temperature and salinity) for a very large fraction of the same set of strains (Opulente et al. 2018, Opulente et al. 2023). The availability of a comprehensive dataset that captures the vast genomic, environmental, and metabolic diversity of yeasts provides a unique testbed for understanding how adaptation to unique environments occurs in eukaryotic genomes (Hittinger et al. 2015).

Several of the pathways that allow yeasts to grow on certain sources are well-characterized (Riley et al 2016). For example, sucrose assimilation depends on the invertase Suc2p, and maltose assimilation depends on the maltose permease Mal31p and maltase (α-D-glucosidase) Mal32, which can also act on sucrose (Ostergaard et al. 2000, Brown et al. 2010). Arguably the best studied pathway is the Leloir or *<u>GAL</u>*actose utilization pathway (Fig. 1), which has become a model not only for understanding gene regulation in eukaryotes (Ptashne & Gann 2001, Johnston 1987), but also for how evolutionary changes in gene sequences, arrangement, and regulation contribute to ecological adaptation (Harrison et al. 2022, Sun et al. 2023, Haase et al. 2021, Venkatesh et al. 2021, Boocock et al. 2021, Hittinger et al. 2010, Slot & Rokas 2010). In the *GAL* pathway of the baker’s yeast *Saccharomyces cerevisiae*, Gal2p or an Hxt transporter protein imports D-galactose into the cell, where the mutarotase domain of Gal10p acts on the sugar, if necessary. Then, Gal1p converts it to galactose-1-phosphate, representing the first energy-consuming step of the pathway (Sellick et al. 2008). Gal7p then converts galactose-1-phosphate to UDP-galactose. Gal10p acts on UDP-galactose using its epimerase domain, resulting in the production of UDP-glucose. Finally, Gal7p converts UDP-glucose to glucose-1-phosphate, which Pgm1p/Pgm2p then converts to glucose-6-phosphate, which enters glycolysis to produce energy for the cell (Sellick et al. 2008).

**Figure 1.**
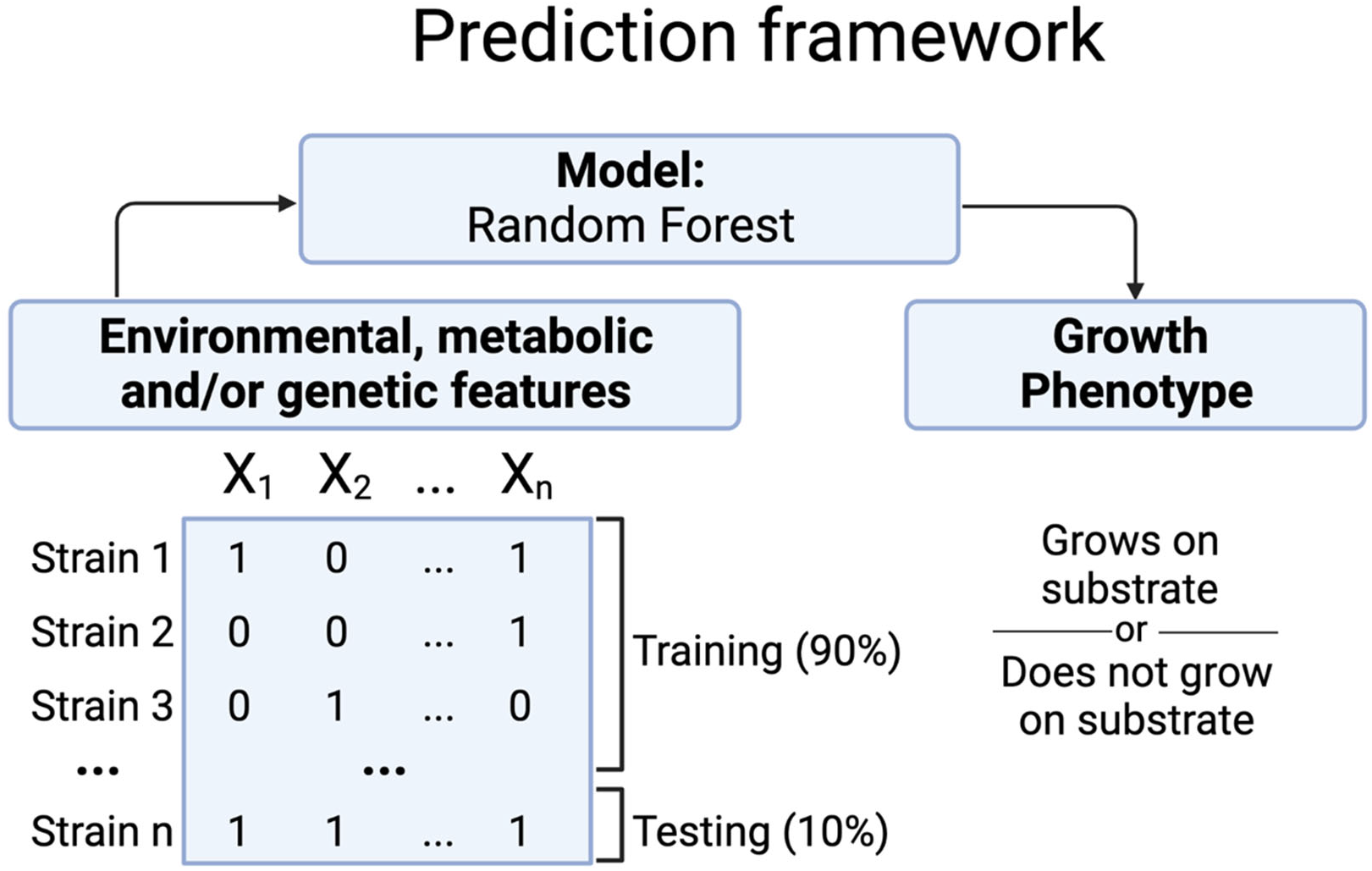
Workflow for machine learning prediction of how diet influences the evolution of primary metabolism in the subphylum Saccharomycotina. Using the phenotype of “grows on substrate” or “does not grow on substrate” for each yeast strain, we trained an XGBoost random forest algorithm on 90% of environmental, qualitative trait, and/or genetic features (893 strains containing 885 species). Using the 10% of remaining data, we tested model performance by looking at accuracy, confusion matrixes, and ROC-AUC curves, and we repeated this assessment 9 more times using cross-validation. Feature importance was calculated using Gini importance as automatically generated by the XGBoost random forest algorithm. Created with BioRender.com.

Galactose abundance varies widely across yeast environments. For example, due to both the dietary influx of galactose and the synthesis of the sugar, galactose is abundant in the gut, bloodstream, and urine of most mammals (including humans) in the form of oligosaccharides, glycoproteins, and glycolipids, as well as in milk and other dairy products in the form of lactose (a disaccharide composed of galactose and glucose subunits) (Brown et al. 2009). Galactose is also found in a variety of fruit, vegetable, and other plant products, such as legumes; levels of galactose in common fruits and vegetables range from <0.1 mg / 100 g to 34 mg / 100 g (Gross & Acosta 1991, Acosta & Gross 1995, Marsilio et al. 2001). Galactose is also part of oligosaccharides, such as lactose, raffinose, and melibiose, as well as glycoproteins and glycolipids, that vary in their distribution across environments (Acosta & Gross 1995, Marsillo et al. 2001); hydrolysis of these molecules by microbial enzymes can release free galactose.

The substantial variation in abundance of galactose in different environments is reflected in the evolution of the *GAL* pathway and its regulation across the subphylum Saccharomycotina. Numerous instances of wholesale pathway loss and gain, including by horizontal gene transfer, have been discovered (Slot & Rokas 2010, Haase et al. 2021, Harrison et al. 2022, Venkatesh et al. 2021), as well as striking instances of ancient, multi-locus polymorphisms within species (Venkatesh et al. 2021, Hittinger et al. 2010, Boocock et al. 2021). Different regulatory systems that lead to different modes of induction and rates of growth have also evolved in different lineages. For example, *C. albicans* exhibits an earlier graded induction in response to galactose, while *S. cerevisiae* has a more bimodal expression (Dalal et al. 2016, Ricci-Tam et al. 2021, Sun et al. 2023).

The rich genomic, environmental, and metabolic data of the Y1000+ Project, coupled with extensive genetic and biochemical knowledge of yeast primary metabolism, provide a unique opportunity to explore the evolution of the genotype-phenotype map, which models the interaction between an organism’s genes and its traits, across a subphylum. However, the enormity and complexity of the Y1000+ Project’s data make standard statistical analyses less suitable. In recent years, machine learning tools, such as support vector machines, random forest algorithms, and convolutional neural networks, have emerged as powerful tools for analyzing biological big data (Zou et al. 2019). Examples include predicting genes involved in specialized metabolism (Moore et al. 2019), predicting protein expression and function from regulatory and protein sequences (Zirmec et al. 2020, Capra et al. 2009, Ma et al. 2018), and distinguishing fungal ecological lifestyles, such as saprobes from plant pathogens (Haridas et al. 2020) or generalists from specialists (Opulente et al. 2023).

In this study, we used a random forest algorithm trained on environmental, metabolic, and/or genomic data to predict growth of nearly all known species of Saccharomycotina on different carbon sources. Predicting growth on 29 different carbon sources tended to be highly accurate when the algorithm was trained on gene presence/absence and/or on presence/absence of growth on other carbon sources, which shows that both metabolic genes and the structure of the metabolic network are highly informative for understanding the evolution of yeast primary metabolism; in contrast, the predictive ability of isolation environment data was weak. Although the most important features associated with prediction accuracy were well-known genes and carbon sources associated with the source of interest, our machine learning approach also identified novel features not previously known to be associated with growth on a given carbon source. To illustrate the predictive ability of our approach, we used growth on galactose as a test case because our machine learning approach suggested a possible novel alternative pathway for galactose assimilation in the genus *Ogataea* and a clade containing *C. auris*, which both lack *GAL* genes. Growth and biochemical assays validated that these species assimilate galactose through a hypothesized oxidoreductive D-galactose pathway, demonstrating the power of machine learning analysis for studying the relationship between genomic and phenotypic variation across vast evolutionary timescales.

## Methods

### Genomic data matrix

Using the KEGG (Kanehisa & Goto 2000, Kanehisa et al. 2023) and InterProScan (Jones et al. 2014) gene functional annotations generated by the Y1000+ Project (Opulente et al. 2023), a data matrix was built with presence and absence of each unique KEGG Ortholog (KO) and counts of each unique InterPro ID number in each genome. Each genome was its own row, and each unique KO (*N =* 5,043) or InterPro ID (*N =* 12,242) present in one or more of the 1,154 yeast genomes was its own column. A python script recorded the presence and absence of KO annotations (Table S1), the number of each InterPro ID for each genome (Table S2), and put them in the appropriate cells of the data matrix. Upon observing that accuracy was typically similar for predicting growth on 29 carbon sources between a random forest algorithm trained just on the KO dataset and the combined KO and InterPro dataset, the KO genomic dataset was used for all subsequent analyses, and the InterPro data was dropped from the genomic analyses following Figure 2. Comparison of our own *GAL* gene searches with the KO dataset revealed that *GAL1* was misannotated, and that the mutarotase and epimerase domains of *GAL10* were annotated separately by KEGG.

**Figure 2.**
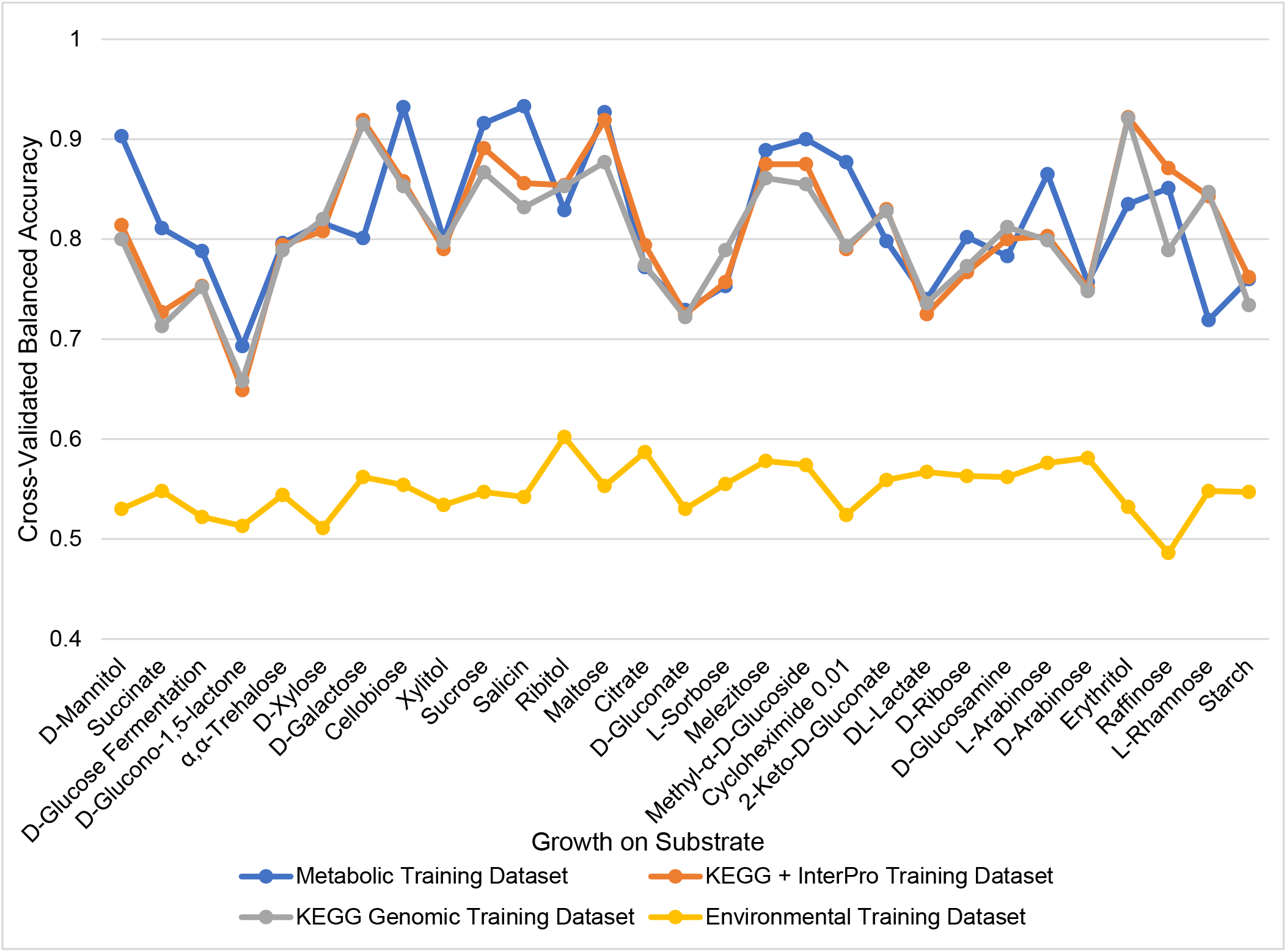
Prediction accuracy of growth on different substrates was high when the random forest algorithm was trained on metabolic data (blue) or genomic data (orange and grey) but low when the algorithm was trained on isolation environment data (yellow). Note that data on growth (and, where applicable, on fermentation) of the condition tested were removed prior to each analysis (e.g., prediction of growth on xylose from metabolic data was conducted using data for growth on all other substrates, but it excluded data for growth on xylose and xylose fermentation). Balanced accuracy was assessed by RepeatedStratifiedKFold(n_splits=10, n_repeats=3) after training the random forest algorithm on either the remainder of the metabolic data, the InterPro and/or KEGG genomic data matrices, or the environmental data. Traits are ordered from most frequent to least frequent in the dataset from left to right. The most important feature for each random forest algorithm, as well as the precision of the algorithm, is shown in Supplementary Table 1.

### Metabolic data matrix

Our metabolic data matrix contained 122 traits from 893 yeast strains from 885 species in the subphylum. The list of traits included growth on different carbon and nitrogen sources, such as galactose, raffinose, and urea, as well as on environmental conditions, such as growth at different temperatures and salt concentrations (Table S3). The metabolic data were sourced from information available for each of the sequenced strains from the CBS strain database. These data were gathered from strains studied as part of the in the published descriptions of species, additional data on strains obtained by previous studies done in the Westerdijk Fungal Biodiversity Institute (CBS), or additional data provided by the depositors of the strains in the CBS culture collection. The data matrix contained metabolic data for 893 / 1,154 species. The percentage of missing data in the data matrix was 37.5% (40,906 missing values out of 108,946 total). Less thoroughly studied traits tended to have more missing data than more commonly found and/or thoroughly studied traits. For example, our data matrix included data on melibiose fermentation, which was estimated to be present in 12% (28/234) of yeasts, but only 26.2% (234/893 of strains have been tested for growth on this substrate. In contrast, our data matrix included data on galactose assimilation, which was estimated to be present in 64.2% (558/868), but 97.2% (868/893) of strains have been tested. Since there were 25 strains for which growth on galactose was not characterized, the total number of strains for which we have both genomic data and galactose assimilation data was 868.

### Environmental data matrix and ontology

The isolation environments for 1,088 (94%) out of the 1,154 yeasts examined were gathered from strain databases, species descriptions, or from *The Yeasts: A Taxonomic Study* (Opulente et al. 2023, Kurtzman et al. 2011). Strains without isolation environments either had been significantly domesticated via crossing or subculturing or were lacking information in our searches. Written descriptions of the environments were converted into a hierarchical trait matrix using a controlled vocabulary. The ontology was built with Web Protégé (https://webprotege.stanford.edu/), with two broader categories: Organic (Fungi, Soil, Animal, and Plant) and Inorganic (Industrial and Soil). Within these categories, more specific controlled vocabulary annotations were connected to each strain: for example, an isolation environment reported as “*Drosophila hibisci* on *Hibiscus heterophyllus*” was associated in our ontology with the animal subclass “*Drosophila hibisci*” and the plant subclass “*Hibiscus heterophyllus*”. This ontology was converted to a binary trait matrix containing all the unique environmental descriptors (Table S4). The same ontology was used in the recent Y1000+ manuscript (Opulente et al. 2023), but that manuscript only considered the first subclass in subsequent analyses; our analyses here used all connections in the ontology for training a random forest algorithm.

### Predicting growth on different carbon sources using machine learning algorithms trained on genomic, metabolic, and / or environmental data

To test whether we could predict growth on 29 different carbon sources from genomic, environmental, and / or (the rest of the) metabolic data, we used a random forest algorithm. These 29 traits were selected because they were measured in at least 743 strains and were present in 20%-80% of strains included in this analysis. For each trait, a random forest algorithm was trained separately on environmental, metabolic, or genomic datasets to evaluate the accuracy of prediction and identify the most important predictive features (Table S5).

We trained a machine learning algorithm built by an XGBoost (1.7.3) random forest classifier (XGBRFClassifier()) with the parameters “ max_depth=12, n_estimators=100, use_label_encoder =False, eval_metric=’mlogloss’, n_jobs = 8” on 90% of the data, and used the remaining 10% for cross-validation, using RepeatedStratifiedKFold from sklearn.model_selection (1.2.1) (Chen & Guestrin 2016, Pedregosa et al. 2011). We used RepeatedStratifiedKFold to generate accuracy measures (including balanced accuracy) and Receiver Operator Characteristic (ROC)/Area Under Curve (AUC) curves for each prediction analysis. We used the cross_val_predict() function from Sci-Kit Learn to generate the confusion matrixes; these matrices show the numbers of strains correctly predicted to grow or not grow on a specific carbon source (True Positives and True Negatives, respectively) and incorrectly predicted (False Positives, predicted to grow but do not; and False Negatives, not predicted to grow but do). Top features were automatically generated by the XGBRFClassifier using Gini importance, which uses node impurity (the amount of variance in growth on a given carbon source for strains that either have or do not have this trait/feature).

In each prediction analysis, we note that we excluded from each training dataset growth and fermentation data for each of the 29 carbon sources under investigation. For example, we excluded growth on galactose and galactose fermentation from the training dataset for predicting growth on galactose; thus, the final metabolic data matrix used in the training contained data from 120 sources and conditions, instead of the total 122. Similarly, we excluded growth on sucrose and sucrose fermentation from the training dataset for predicting growth on sucrose; we excluded xylose and xylose fermentation from the training dataset for predicting growth on xylose.

### *GAL1*, *GAL7*, *GAL10*, and *GAL102* gene searches

To determine presence / absence of genes in the *GAL* pathway in each of the genomes of the 1,154 strains included in our study, we conducted sequence similarity searches for the *GAL1*, *GAL7*, *GAL102*, and *GAL10* genes using the jackhmmer function from the HMMER software, version 3.3.2 (Eddy 2009). Using the representative *GAL* gene sequences from the *Candida albicans* genome, jackhmmer searched for all hits above a similarity score of 200, which captured genes from all 12 Saccharomycotina taxonomic orders, and then used these results to build a new profile to search for the gene throughout the phylogeny. jackhmmer repeated this method until the results converged, which was three rounds for all genes except *GAL10*, which required five rounds, likely because the mutarotase and epimerase domains are part of the same protein in some yeast orders (e.g., Saccharomycetales and Serinales) but belong to two separate proteins (encoded by *GALM* and *GALE*, respectively) in others (e.g., Lipomycetales) (Haase et al. 2021, Slot & Rokas, 2010). In analyses where only the *GAL* gene dataset was used as genomic data, both the presence / absence and similarity score produced by jackhammer for *GAL1*, *GAL7*, and *GAL10* were included in the dataset; hits with similarity scores below 200 were considered absent and were entered as 0 (Table S6). As noted above, comparison of our own *GAL* gene searches with the KO dataset revealed that *GAL1* was misannotated, and that the mutarotase and epimerase domains of *GAL10* were annotated separately by KEGG.

### Quantification of galactose utilization in strains lacking the *GAL* pathway

To validate galactose utilization by certain strains lacking the GAL genes that were identified in our qualitative metabolic data matrix, we quantified growth and galactose consumption in liquid culture. Standard undefined yeast lab media was prepared as previously described (Sherman 2002). YPD medium for culturing yeasts contained 10 g/L yeast extract, 20 g/L peptone, 20 g/L glucose, and 18 g/L agar (US Biological). Cells were streaked onto YPD plates, and single colonies were picked. Cells were inoculated into 5 mL of YP (10 g/L yeast extract, 20 g/L peptone) + 2% galactose (Amresco) and grown to mid-log phase (48 – 55 hours depending on the strain, see Table S10 for further information) on a tissue culture wheel at room temperature.

The optical density of the cells was measured at 600 nm (OD_600_) using an OD600 DiluPhotometer (Implen). Cells were inoculated into 50 mL YP + 2% galactose at a starting OD_600_ 0.05 for all species except for the negative control species, *Saccharomycopsis malanga*, which was inoculated at starting OD_600_ 0.01 due to the low cell density caused by the absence of its *GAL* pathway. The cultures were shaken in non-baffled 150-mL Erlenmeyer flasks (Fisher Scientific) at 250 rpm at room temperature for seven days. 1 mL of culture was collected every 24 hours and spun down; 600 µL of supernatant were used for extracellular sugar quantification via high performance liquid chromatography and refractive index detection (HPLC-RID). OD_600_ readings were also taken at each 24-hour timepoint. All samples taken for HPLC-RID were stored at -20 °C until the end of the experiment. Extracellular galactose concentrations were determined by HPLC-RID as previously described using a galactose standard (Schwalbach et al. 2012, Lee et al. 2021). The strain *S. cerevisiae gre3Δ::loxP-kanMX-loxP* (Parreiras et al. 2014) served as a positive control for galactose utilization because it has an intact *GAL* pathway; the deletion of *GRE3*, which encodes a promiscuous aldose reductase that could conceivably have some activity on galactose (Masuda et al. 2008), also allowed this strain to serve as a negative control for the hypothesized oxidoreductive pathway. Galactose concentrations were expressed as g/L, and the results correspond to the mean value of biological triplicate timepoints. All extracellular galactose quantification data visualization was performed using R (v4.1.2) in the RStudio platform (v2022.07.01+554) and with the package ggplot2 (v3.4.2) (R Core Team 2021, RStudio Team 2022, Wickham 2016).

### Assay for galactose- and NADPH-dependent enzymatic activity

To determine whether galactose utilization in strains lacking the *GAL* genes but able to grow in galactose occurred through a hypothesized oxidoreductive D-galactose pathway, we tested NADPH-dependent enzymatic activity on galactose as a sole carbon source. Yeast cells were pregrown in YPD, single colonies were inoculated into 5 mL YP + 2% galactose, cultures were grown to mid-log phase, and they were inoculated into 50 mL YP + 2% galactose using the same methods as described above. *Candida duobushaemulonii*, *Candida ruelliae*, and *Ogataea methanolica* cells were harvested at mid-log phase along with their respective *S. cerevisiae gre3Δ::loxP-kanMX-loxP* negative controls for whole-cell lysate protein extraction using Y-PER (Thermo Fisher Scientific). 1 mL of culture was sampled, and cells were centrifuged at 3,000 x *g* at 4 °C for 5 minutes. 250 mg of wet cell pellet were resuspended in 1,250 µL of Y-PER and homogenized by pipetting. The mixture was left to agitate at room temperature for 50 minutes to ensure successful cell lysis and soluble protein extraction. Cell debris was pelleted at 14,000 x *g* for 10 minutes at room temperature. Finally, 1 mL of supernatant was removed for analysis and protein concentration determination. Protein concentrations were determined using the Pierce BCA protein assay kit and protocol (Pierce Biotechnology), and absorbance at 562 nm was measured using The Infinite M1000 microplate reader (Tecan). Galactose-dependent enzymatic activity was determined by monitoring the oxidation of the cofactor NADPH to NADP^+^ by absorbance measurement at 340 nm at 25 °C (Cadete et al. 2016). The assay mixture (200 µL) contained 200 mM Tris-HCl (pH 7.5), 5 mM of NADPH, 200 mM of galactose, 200 µg of undefined cell-free protein extract, and deionized water in 96-well plates (Corning 96 Well Clear Flat Bottom UV-Transparent). In addition, each assay contained a protein extract blank and a substrate (without galactose) blank to account for protein and substrate noise, cofactor degradation, and off-target cofactor oxidation. Enzyme assays were performed in biological quadruplicate. Data analyses and plots were performed and visualized using the methods described above.

## Results

### Machine learning accurately predicts growth on 29 different carbon sources from metabolic and genomic data but not from environmental data

A random forest algorithm (Figure 1) trained on the metabolic data matrix had high balanced accuracy (on average, 82%) for predicting growth of the 893 strains representing 885 of the Y1000+ yeast species on 29 different carbon sources. This result indicates that variation in the content and structure of the primary metabolic network in different strains informs patterns of growth on these substrates (Figure 2, Table S5). A random forest algorithm trained on the genomic data matrices (comprised of InterPro and/or KEGG Orthology (KO) annotations) was similarly accurate for predicting growth on these 29 sources (on average, 80-81% balanced accuracy). In contrast, when the random forest algorithm was trained on environmental datasets, the balanced accuracy was between 48-61% (on average, 55%), which is only marginally above random accuracy (Figure 2, Table S5). This result suggests that our environmental dataset does not provide useful predictors for growth on these sources. Examination of the ROC/AUC curves, confusion matrixes, and most important features for predicting growth on xylose, sucrose, and galactose supports this hypothesis: accuracy is only marginally above random using environmental data, and the most important features concern isolation environments not known to have high amounts of these sugars (Figure S1).

### Top features for predicting growth on a specific carbon source are related sources and metabolic genes

The top features for predicting growth on the 29 carbon sources examined were often biologically relevant (Figure 2, Figure 3, Table S5). For example, for xylose, the most important feature was growth on xylitol, a metabolic intermediate in the typical xylose-degrading pathway in yeasts and other fungi (Wohlbach et al. 2011, Meng et al. 2022), while for sucrose, the most important feature was maltose, another disaccharide containing a glucose moiety (Brown et al. 2011) (Figure 3). For galactose, the top features included 2-keto-D-gluconate and L-sorbose, which are generated from glucose or galactose, respectively, by the enzymes acting on an alternative galactose-degrading pathways in some bacteria and fungi (Fekete et al. 2004, Tanimura et al. 2003, Meng et al. 2022, Sun et al. 2020), as well as lactose and melibiose, polysaccharides that contain galactose (Figure 3).

**Figure 3.**
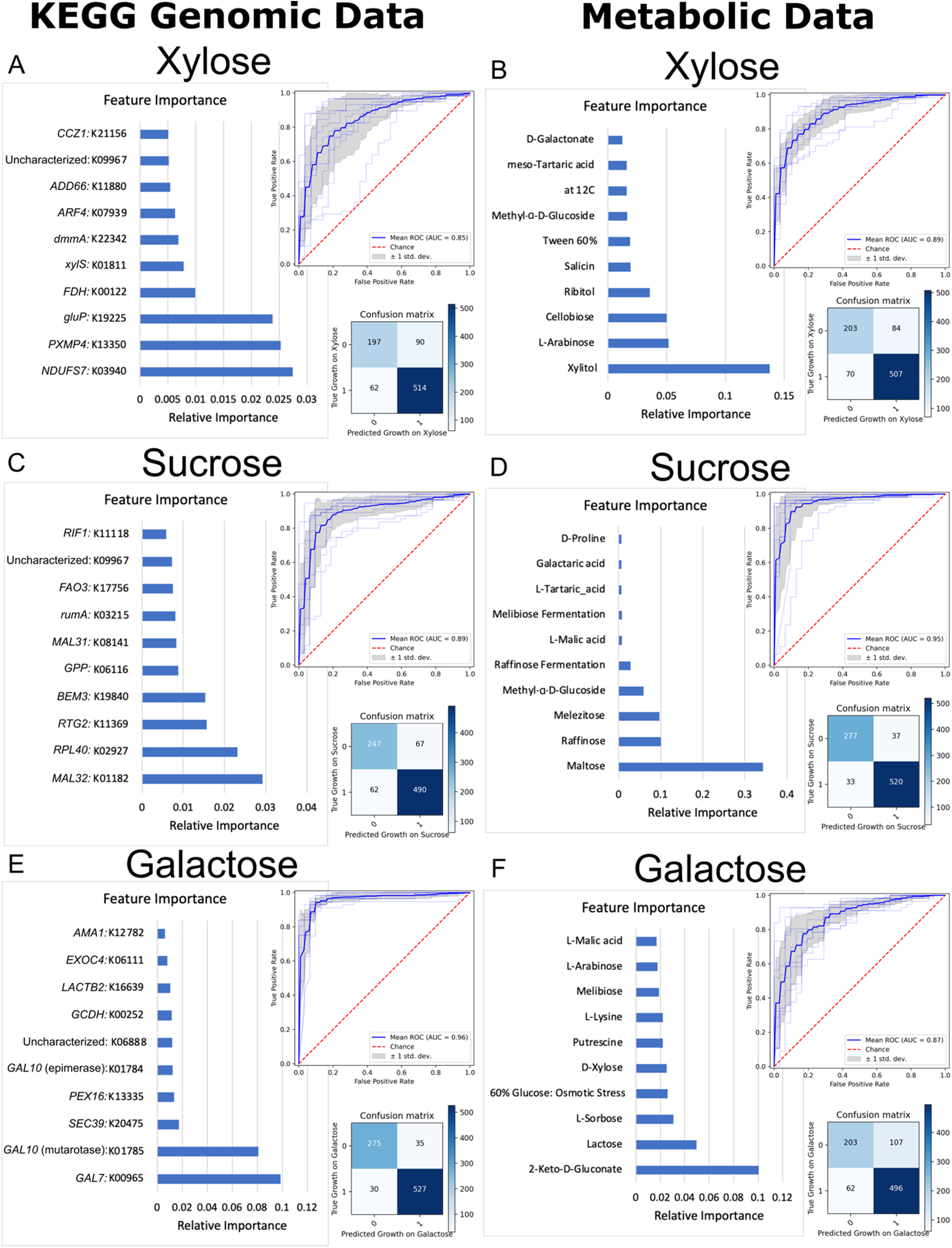
Prediction accuracy of growth on different sugars was high when the random forest algorithm was trained on genomic data (A, C, E), and similarly high when the algorithm was trained on metabolic data (B, D, F). Panels A and B: prediction of growth on xylose from genomic (A) or metabolic data (B). Panels C and D: prediction of growth on sucrose from genomic (C) or metabolic (D) data. Panels E and F: prediction of growth on galactose from genomic (E) or metabolic (F) data. Note that data on growth (and, where applicable, on fermentation) of the carbon source tested were removed prior to each analysis (e.g., prediction of growth on xylose from metabolic data was conducted using data for growth on all other substrates and conditions, but it excluded data for growth on xylose and xylose fermentation). Also note that KEGG Ontology misannotated *GAL1,* likely leading *GAL1* to not be in the top features, and that the epimerase and mutarotase domains encoded by *GAL10* were annotated separately by this program. Accuracy is shown in the form of confusion matrices, which show strains predicted correctly to not grow on the sugar (true negatives, top left), strains predicted to grow on the sugar that do not (false positives, top right), strains correctly predicted to grow on the sugar (true positives, bottom right), and strains predicted to not grow on the sugar that do (false negatives, bottom left), as well as Receiver Operating Characteristic (ROC) curves, which show the true positive rate over false positive rate with changing classification thresholds. Feature importance graphs are also included to show the input features that are most useful for predicting growth on this sugar. XGBoost random forest was used to generate feature importance, and cross_val_predict() from sklearn.model_selection was used to generate confusion matrices. ROC curves were generated using the roc_curve function from sklearn.metrics. The prediction accuracies of growth on xylose, sucrose, and galactose from isolation environment data are shown in Supplemental Figure 1.

**Figure 4.**
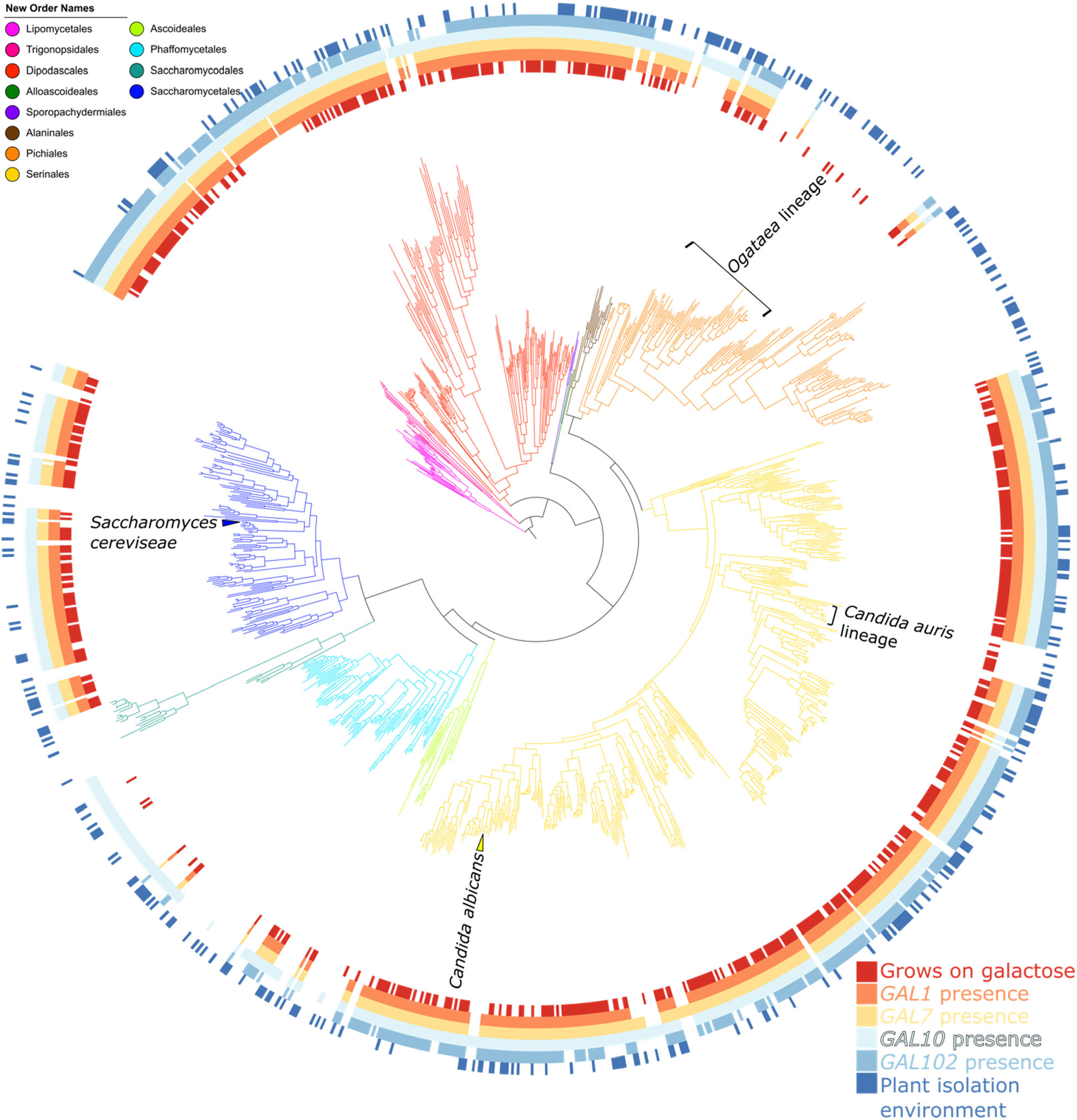
Distribution of *GAL* genes and plant isolation environments across the yeast phylogeny. The ability of the different strains to grow on galactose, the presence of genes *GAL1*, *GAL7*, *GAL10*, and *GAL102*, and whether they were isolated from plant environments are plotted as circles around the yeast phylogeny. Strain names are omitted for easier visualization, but they can be found in Figure S2. The colors of the different branches of the phylogeny correspond to the 12 taxonomic orders (Groenewald et al. 2023).

A random forest algorithm trained on KEGG Orthology (KO) annotations was similarly accurate for predicting growth on xylose (85%), sucrose (88%), and galactose (92%) to the combined KO and InterPro genomic dataset (Figure 2, Figure 3). Despite the larger size of the genomic data matrix (over 5,000 features compared to the metabolic data matrix of 122 features), the top features of the genomic data matrix were still often related to genetic pathways or enzymes known to be involved in the utilization of each source. The top features for the highly accurate prediction of growth on galactose were *GAL7* and *GAL10* (specifically the mutarotase domain), which are parts of the yeast *GAL* pathway (Harrison et al. 2022). Despite the mis-annotation of the yeast *GAL1* by KO (see Methods), the algorithm was still nearly as accurate when trained on the entire genomic data matrix as when trained on the manually curated *GAL* gene orthologs (Figure 5). The top feature for the algorithm predicting growth on sucrose was oligo-1,6-glucosidase (K01182), which corresponds to the α-glucosidases encoded by *MAL32* and *MAL12*, as well as *IMA1-IMA5*, which indeed do act on sucrose, as well as maltose in some yeasts (Brown et al. 2010, Ostergaard et al. 2000). The distribution of *XYL1, XYL2,* and *XYL3* does not always correlate with yeast growth on xylose (Riley et al. 2016, Nalabothu et al. 2023). Even though the *XYL* genes were present in the KO database (with the exception of *XYL3*, which was misannotated), they were not among the top features contributing to the 85% prediction accuracy, but an α-xylosidase (K01811) was the fifth most important feature (Figure 3). Since galactose metabolism and its associated genetic pathway has been thoroughly studied in yeasts, the remainder of this paper is focused on using growth on galactose as a test case for the utility of this machine-learning pipeline.

**Figure 5.**
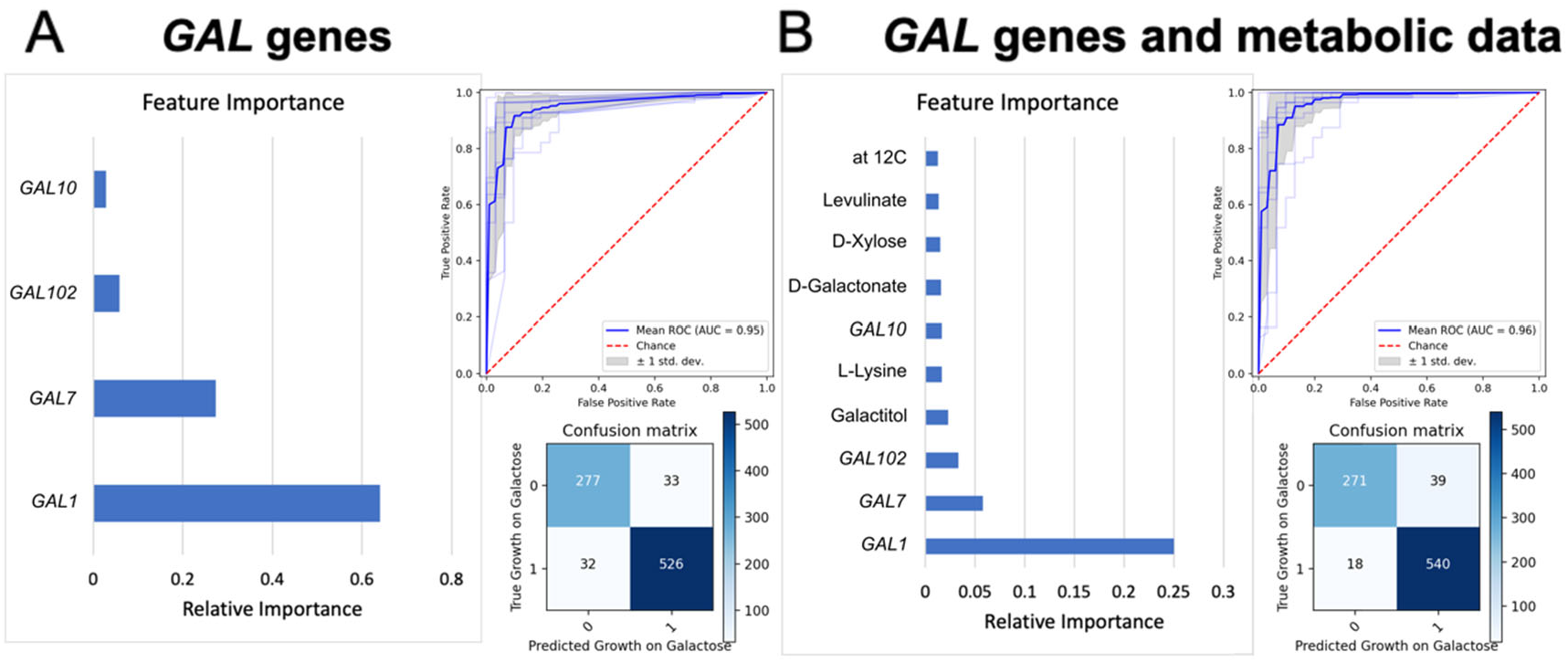
*GAL* gene presence / absence and ability to grow on galactitol are highly predictive of growth on galactose across the subphylum Saccharomycotina. **A.** Using the presence / absence patterns of the genes *GAL1, GAL7, GAL10,* and *GAL102* as input data, the XGBoost random forest algorithm predicted growth on galactose with high accuracy, as shown by the confusion matrix, the ROC/AUC curve, and the individual feature importance. **B.** Using both the presence / absence patterns of *GAL* genes (from panel A) and metabolic data, the algorithm predicted growth on galactose with even higher accuracy, shown by the confusion matrix, the ROC/AUC curve, and the individual feature importance. Note that, after *GAL1*, *GAL7*, and *GAL102* genes, growth on galactitol is the next most important feature for predicting growth on galactose.

### The GAL genes are highly predictive of growth on galactose in most, but not all, strains

Plotting the presence / absence of the *GAL* genes jointly with the presence / absence of growth on galactose on genome-scale phylogeny of 1,154 yeast strains showed that the distributions of the *GAL* genes were tightly correlated with the distribution of growth on galactose. Specifically, 526 / 558 strains that can grow on galactose have the *GAL* genes, and 277 / 310 strains that cannot grow on galactose lack the *GAL* genes. Notably, there are two lineages in the orders Serinales and Pichiales that can grow on galactose but lack the *GAL* genes (Figure 4). One lineage contains species closely related to the emerging opportunistic pathogen *Candida auris* in the order Serinales. The second lineage contains species belonging to the genus *Ogataea* in the order Pichiales. Isolation environments, such as isolation from plants, showed no significant association with growth on galactose (Figure 4).

Using the scores from the sequence similarity searches (from the jackhmmer software) of *GAL1, GAL7*, *GAL102,* and *GAL10*, growth on galactose was even more accurately predicted, at 92.6%. When the metabolic dataset was added to the training data, the accuracy increased even further to 93.4% (Figure 5). This increase in accuracy suggests that there are strains for which presence or absence of the *GAL* genes cannot accurately predict growth on galactose; if that were the case, then the increase in accuracy due to the inclusion of the rest of the metabolic dataset raises the possibility that there might be an alternative galactose-degrading pathway in some yeasts. After the *GAL* genes, the most predictive feature was growth on galactitol, pointing to a possible role for this metabolite as an intermediate in a potential alternative pathway (Figure 5). Previous work in filamentous fungi identified a galactose-degrading pathway that involves galactitol as an intermediate (Chroumpi et al. 2022, Fekete et al. 2004), leading us to hypothesize that a similar pathway may be present in these yeasts and contribute to the increase in accuracy.

### Machine learning predicts an alternative galactose-degrading pathway in two yeast lineages that lack *GAL* genes

To further explore the possibility of an alternative galactose utilization pathway that uses galactitol as an intermediate, we trained our random forest algorithm just on the *GAL* genes and growth on galactitol. We found that this algorithm was just as accurate as when the rest of the metabolic dataset was added (93.6% versus 93.4%). Examination of the confusion matrices when the algorithm was trained using just the *GAL* gene data versus when trained on the *GAL* gene data and metabolic data suggested that the increase in accuracy came from 15 species that were previously classified as false negatives and were now true positives (Figure 6). Since these species lack the *GAL* genes, our original algorithm predicted that they could not grow on galactose; when growth on galactitol was added, however, they were correctly predicted to grow on galactose, further supporting the hypothesis that they have an alternative galactose-degrading pathway (Figure 6). These 15 species are all able to grow on galactitol and belong to the two lineages that lack *GAL* genes, as noted previously in Figure 4: the lineage of species closely related to *Candida auris* in Serinales and the genus *Ogataea* in Pichiales. Even with this highly accurate algorithm, there were several species that were still not correctly predicted: 18 false negatives (strains that are predicted not to grow, but do) (Table S7) and 39 false positives (strains that are predicted to grow, but do not) remained (Table S8). These species warrant further investigation as they may contain other alternative pathways, grow weakly on galactose or only under specific conditions, use galactose in glycosylation but not for assimilation (as the fission yeast *Schizosaccharomyces pombe*) (Suzuki et al. 2010), or have pseudogenized *GAL* genes (Hittinger et al. 2004). We note that the *GAL* genes of yeasts that were false positives in our classification exhibited, on average, lower sequence similarity scores in our *GAL* gene searches than the *GAL* genes of yeasts that were true positives (Table S9), which is consistent with reduced purifying selection.

**Figure 6.**
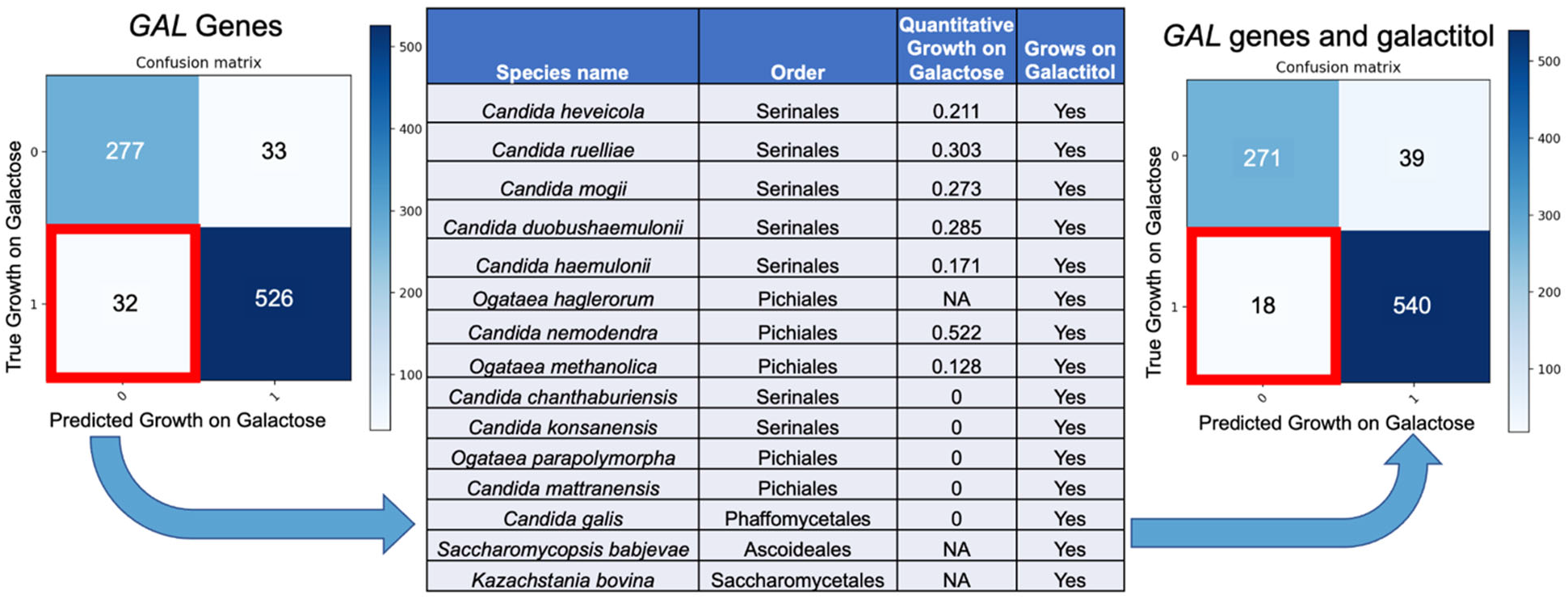
Adding the galactitol growth data to presence / absence of the *GAL* genes increased prediction accuracy by correctly classifying several false negatives as true positives. On the left is the confusion matrix for predicting growth on galactose using just *GAL1*, *GAL7, GAL10*, and *GAL102* presence / absence. Note the presence of 32 false negatives; the algorithm predicted that these 32 species would be unable to grow on galactose because they lack the *GAL* genes, but they are known to grow on galactose. When the metabolic trait “Growth on Galactitol” was added to the training data, 15 of these species were now correctly predicted to grow on galactose and were moved to the “True Positive” category, while 17 remained false negatives. One additional species that has low sequence similarity scores for the presence of *GAL* genes in its genome (*Kuraishia hungarica*) also became a new false negative, bringing the total up to 18 false negatives, as shown in the confusion matrix on the right. The taxonomy (order) (Groenewald et al. 2023), quantitative growth on galactose (which is normalized to growth on glucose), and qualitative ability to grow on galactitol for these 15 species are listed in the table.

### Some Pichiales and Serinales species utilize galactose through an oxidoreductive galactose utilization pathway

To test the hypothesis that some species lacking *GAL* pathways can indeed utilize galactose, we tested three species (Table S10) from two different orders, *C. ruelliae* and *C. duobushaemulonii* from Serinales and *O. methanolica* from Pichiales, for growth on galactose as the sole carbon source and measured galactose consumption (Groenewald et al. 2023). All three species grew to high cell densities and accumulated more biomass than the *S. cerevisiae* positive control (Figure S3A), which contains an intact *GAL* pathway. Sugar quantification indicated galactose consumption in all three species (Figure 7A). The first step of the known oxidoreductive galactose pathway utilizes an aldose reductase, which reduces galactose to the sugar alcohol galactitol while oxidizing NADPH to NADP^+^ (Seiboth & Metz 2011) (Figure 7B). Thus, we developed a biochemical assay for NADPH-dependent enzymatic activity on galactose as the sole carbon source. In this assay, species that exhibit the hypothesized enzymatic activity are predicted to show a decrease in NADPH absorbance at 340 nm over time, while species that do not exhibit enzymatic activity are predicted to show no decrease in NADPH over time (Figure 7C). All three species displayed decreases in absorbance of NADPH compared to their respective negative controls with no substrate (Figure 7D) and no extracted protein (Figure S3B), which indicates that the cells express NADPH-dependent enzymatic activity that is dependent on the presence of galactose. The *S. cerevisiae* negative control used for this experiment possessed an intact *GAL* pathway and did not show a decrease in NADPH absorbance over time, indicating a lack of NADPH-dependent enzymatic activity on galactose as the sole carbon source. Thus, we conclude that these three species possess at least the first step of an oxidoreductive pathway.

**Figure 7.**
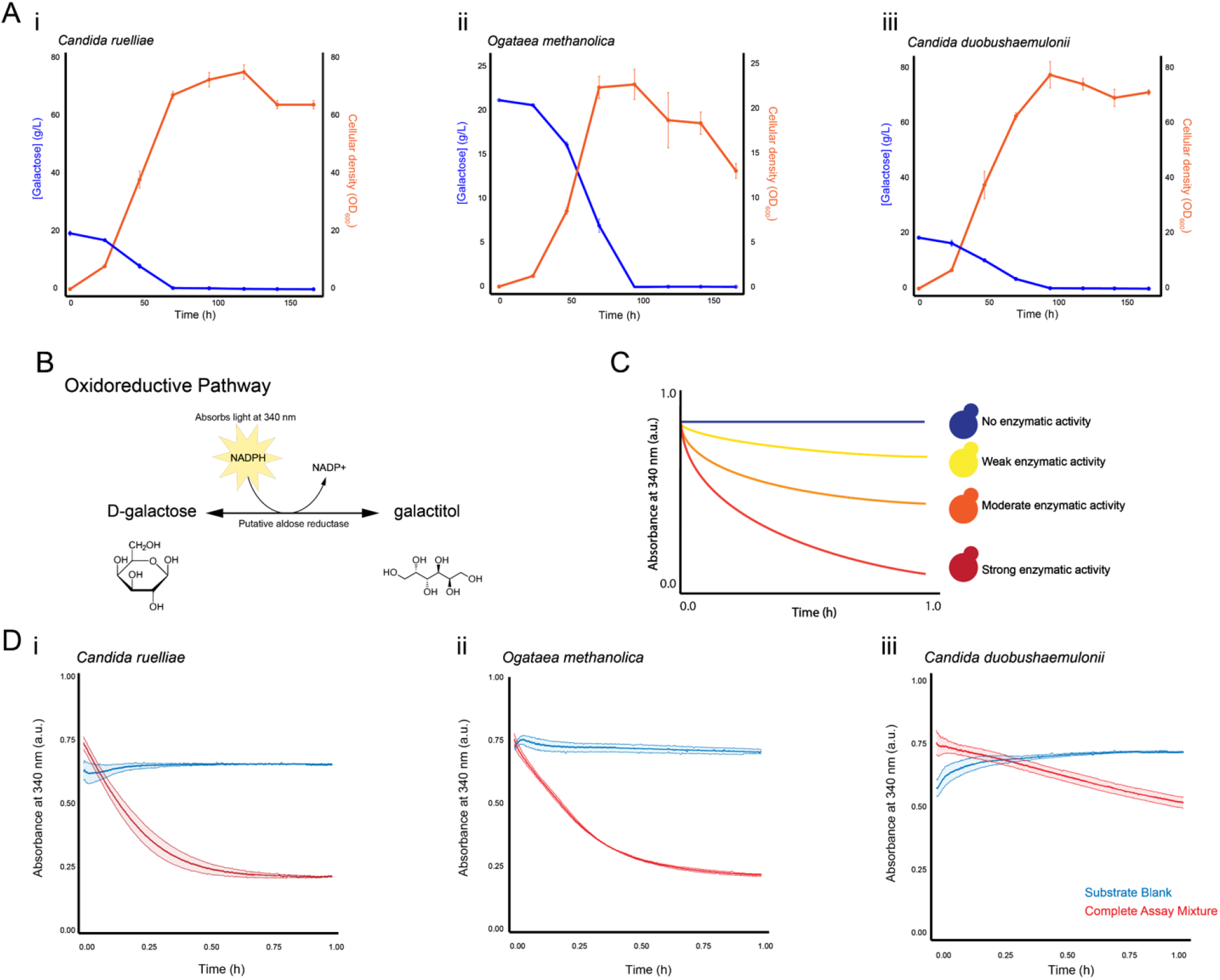
All three species showed galactose consumption and NADPH-dependent enzymatic activity on galactose. A. Average and standard deviation across three biological replicates of galactose concentrations present in medium with galactose as the sole carbon source (blue) and OD_600_ growth measurements (orange) for *C. ruelliae* (i), *O. methanolica* (ii), and *C. duobushaemulonii* (iii) over 168 hours. **B.** Schematic diagram of the first step of a hypothesized oxidoreductive galactose pathway using an aldose reductase to reduce galactose to galactitol by oxidizing NADPH to NADP^+^. **C.** Illustration of the expected results for different levels of enzymatic activity. As the amount of NADPH present in the assay mixture decreases, absorbance at 340 nm decreases. **D.** Average and standard deviation across four biological replicates of NADPH absorbance at 340 nm over time comparing the complete assay mixture (red) to a substrate blank with no galactose added (blue) for *C. ruelliae* (i), *O. methanolica* (ii), and *C. duobushaemulonii* (iii).

## Discussion

In this study, we employed machine learning on the rich environmental, metabolic, and genomic data from nearly all known species of an entire eukaryotic subphylum to predict each strain’s growth on different carbon sources. We found that we could accurately predict growth on diverse sources of carbon from genomic and / or metabolic data but not from environmental data (Figure 2). Previous research showed that many yeast traits are connected in a trait-trait network, likely due to shared genes in different metabolic pathways (Opulente et al. 2018, Opulente et al. 2023). These connections and overlap in gene functions likely explain the high accuracy of prediction from metabolic and / or genomic data. Interestingly, accuracy of prediction was high, even for carbon sources for which enzyme specificity was lacking, such as xylose (Figure 3) (Nalabothu et al. 2023). However, accuracy for xylose growth was lower than for predicting growth on sources, such as galactose, whose utilization pathways contain dedicated enzymes (Figure 3).

In contrast, the accuracy of prediction of growth on different carbon sources from isolation environment data was marginally better than random (Figure 3). There are two possible explanations for this finding. The first is that isolation environments may be heterogenous in their carbon sources, supporting metabolically diverse yeasts. An alternative, not necessarily mutually exclusive explanation, is that isolation environments can be informative with respect to yeast diets, but that our current environmental data are incomplete. Notably, our isolation environmental data for each yeast included in the data matrix stem from information present in the taxonomic description of the type strain of each species. A dataset that contains the range of isolation environments of each yeast species would potentially be much more informative but is currently unavailable.

We also found that machine learning accuracy for predicting growth on galactose was higher when both the presence / absence of *GAL* genes and growth on galactitol were used in training compared to just the presence / absence of the *GAL* genes alone (Figure 5), suggesting the presence of a rare alternative galactose-degrading pathway. We discovered that this alternative galactose-degrading pathway is found in two distinct lineages that grow in galactose in the absence of *GAL* genes; we further proposed that this alternative pathway involves galactitol as a metabolic intermediate (Figures 4-6). Enzyme assays validated the oxidoreductive activity of three species in these two lineages when grown on galactose, providing additional support for the hypothesized mechanism of utilization (Figure 7). We are currently investigating the genes involved in this alternative pathway. This work illustrates the remarkable breadth of yeast metabolic diversity and how machine learning approaches can help uncover novel biology, even in well-studied traits, such as galactose assimilation.

The potential for additional discoveries using machine learning is further highlighted by considering the several yeasts that appear as false positives or false negatives in our machine learning predictions. There are several possible explanations for why we currently cannot accurately predict growth on galactose for every strain in the subphylum. One explanation for some of the false positives could be that the *GAL* pathway is inactivated in some of the strains examined, but that their genomes contain *GAL* pseudogenes. Examples of *GAL* pseudogenes are known from several different species (Hittinger et al. 2004, Hittinger et al. 2010, Venktesh et al. 2021), but strains with pseudogenes would still give positive hits in our ortholog detection analyses. In support of this hypothesis, the average sequence similarity scores for the *GAL* genes in yeasts classified as false positives were lower than the scores for *GAL* genes in yeasts classified as true positives (Tables S8 and S9). Another possible explanation for false positives could be that some yeasts may contain *GAL* genes that are used in other processes, such as glycosylation, but not in assimilation; although such examples are not currently known from the Saccharomycotina, the fission yeast *Schizosaccharomyces pombe* (subphylum Schizosaccharomycotina) is a case in point (Matsuzawa et al. 2011). They may also be growing very weakly or under specific conditions not tested here. Also, since growth on galactitol is predictive of this alternative pathway of galactose utilization, the algorithm now predicts that any strain that grows on galactitol can also grow on galactose, which may not always true (e.g., they may be still missing the gene to convert galactose to galactitol). In fact, there are six yeasts in the list of false positives that grow in galactitol but do not grow in galactose. Finally, since more false positives are in lineages other than the more extensively studied Serinales and Saccharomycetales, fewer strains to train on may lead to less accurate gene characterization by jackhmmer; alternatively, the induction of *GAL* genes or use of the pathway may be different in these lineages(Table S8).

Yeasts that appear as false negatives in our analyses, which indicates that they can indeed grow on galactose but are predicted to not be able to do so by the random forest algorithm, may be growing weakly or they may have other alternative pathways that do not involve galactitol. These may also lack the correct inducing conditions to test positive for growth on galactitol since they are often closely related to our documented alternative pathway species (Table S7). Additionally, seven (out of 18) of these have *GAL* genes that are highly divergent in their sequences, indicating that they may have homologs that do not reach the sequence similarity threshold (Table S7). These yeasts could have very divergent, but still functional, *GAL* genes or they may have been misannotated or have incomplete genomes that are missing the complete version of the *GAL* genes, which would cause a lower sequence similarity score.

These results demonstrate that machine learning, particularly random forests, is a powerful approach to find important traits in genomic and metabolic datasets and for investigating the evolution of the yeast genotype-phenotype map. This tool is likely to prove useful for looking at many different pathways and phenotypes, including non-metabolic ones (e.g., cactophily, cell morphology), in the future.

## Acknowledgements

Thank you to Tony Capra, the Hittinger Lab, the Rokas Lab, and Y1000+ Project team members for helpful discussions throughout the duration of this project; Trey K. Sato for the control strain of *S. cerevisiae*; and Mick McGee and the GLBRC Metabolomics Facility for metabolite quantification. This work was performed using resources contained within the Advanced Computing Center for research and Education at Vanderbilt University in Nashville, TN. X.X.S. was supported by the National Science Foundation for Distinguished Young Scholars of Zhejiang Province (LR23C140001), the Fundamental Research Funds for the Central Universities (226-2023-00021), and the key research project of Zhejiang Lab (2021PE0AC04). This work was supported by the National Science Foundation (grants DEB-2110403 to C.T.H. and DEB-2110404 to A.R.). Research in the Hittinger Lab is also supported by the USDA National Institute of Food and Agriculture (Hatch Project 1020204), in part by the DOE Great Lakes Bioenergy Research Center (DOE BER Office of Science DE–SC0018409, and an H. I. Romnes Faculty Fellowship (Office of the Vice Chancellor for Research and Graduate Education with funding from the Wisconsin Alumni Research Foundation). Research in the Rokas lab is also supported by the National Institutes of Health/National Institute of Allergy and Infectious Diseases (R01 AI153356), and the Burroughs Wellcome Fund.

## Supplementary Figures and Tables

**Supplementary Figure 1. Prediction accuracy of growth on different sugars was low when the random forest algorithm was trained on environmental data.** Prediction of growth on xylose (panel **A**), sucrose (panel **B**), and galactose (panel **C**) from environmental data. The right side of each panel shows the relative importance of different features (feature importance), i.e., the input features that are most useful for predicting growth on a given sugar. The top right graph of each panel is the receiver operating characteristic (ROC) curve, which shows the true positive rate over false positive rate with changing classification thresholds. At the bottom right of each panel is the accuracy of classification in the form of a confusion matrix. Each confusion matrix shows strains predicted correctly to not grow on the sugar (true negatives, top left), strains predicted to grow on the sugar that do not (false positives, top right), strains correctly predicted to grow on the sugar (true positives, bottom right), and strains predicted to not grow on the sugar that do (false negatives, bottom left). Xgboost random forest used to generate feature importance, and cross_val_predict() from sklearn.model_selection used to generate confusion matrixes. ROC curves were generated using the roc_curve function from sklearn.metrics.

**Supplementary Figure 2. Distribution of *GAL* genes and plant isolation environments across the Saccharomycotina yeast phylogeny.** The ability of the different strains to grow on galactose, the presence of genes *GAL1*, *GAL7*, *GAL10*, and *GAL102*, and whether they were isolated from plant environments are plotted as circles (from innermost to outermost) around the Saccharomycotina yeast phylogeny. The colors of the different branches of the Saccharomycotina phylogeny correspond to the 12 taxonomic orders.

**Supplementary Figure 3. Positive and negative control data for experiments in Figure 7.** A. Average and standard deviation across three biological replicates of galactose concentrations in the medium (blue) and OD_600_ growth measurements (orange) for the positive control species *S. cerevisiae* (i) and the negative control species *Saccharomycopsis malanga* (ii). B. Average and standard deviation across four biological replicates of NADPH absorbance at 340 nm over time for the negative control *S. cerevisiae* (red), the substrate blank for the negative control (blue), and protein blank for all species (yellow). The same protein blanks were used for all species included in the enzyme assay since each replicate of the enzyme assay included all four species on one 96-well plate and the protein blank possessed reagents which were the same across all species (Tris-HCl, galactose, NADPH, and deionized water). Note that *S. cerevisiae* is a positive control for growth (Panel A) and a negative control for galactose reductase activity (Panel B).

**Supplementary Table 1. InterPro genomic data matrix.**

**Supplementary Table 2. KEGG Orthology genomic data matrix.**

**Supplementary Table 3. Metabolic data matrix.**

**Supplementary Table 4. Isolation environmental data matrix.**

**Supplementary Table 5. Accuracy, precision, and most important trait of a random forest algorithm trained on metabolic, genomic, and environmental data to predict growth on 29 substrates.**

**Supplementary Table 6. Jackhmmer *GAL* gene sequence similarty score for every strain used to train the random forest algorithm.**

**Supplementary Table 7. Strains classified as false negative for algorithms trained on the *GAL* genes and metabolic data.**

**Supplementary Table 8. Strains classified as false positives for algorithms trained on the *GAL* genes and metabolic data.**

**Supplementary Table 9. Jackhmmer *GAL* gene sequence similarity scores for false positives were on average lower than correctly classified strains.**

**Supplementary Table 10. All of the species used for functional experiments.**

