## Supplementary Figures for "Machine learning illuminates how diet influences the evolution of yeast galactose metabolism"

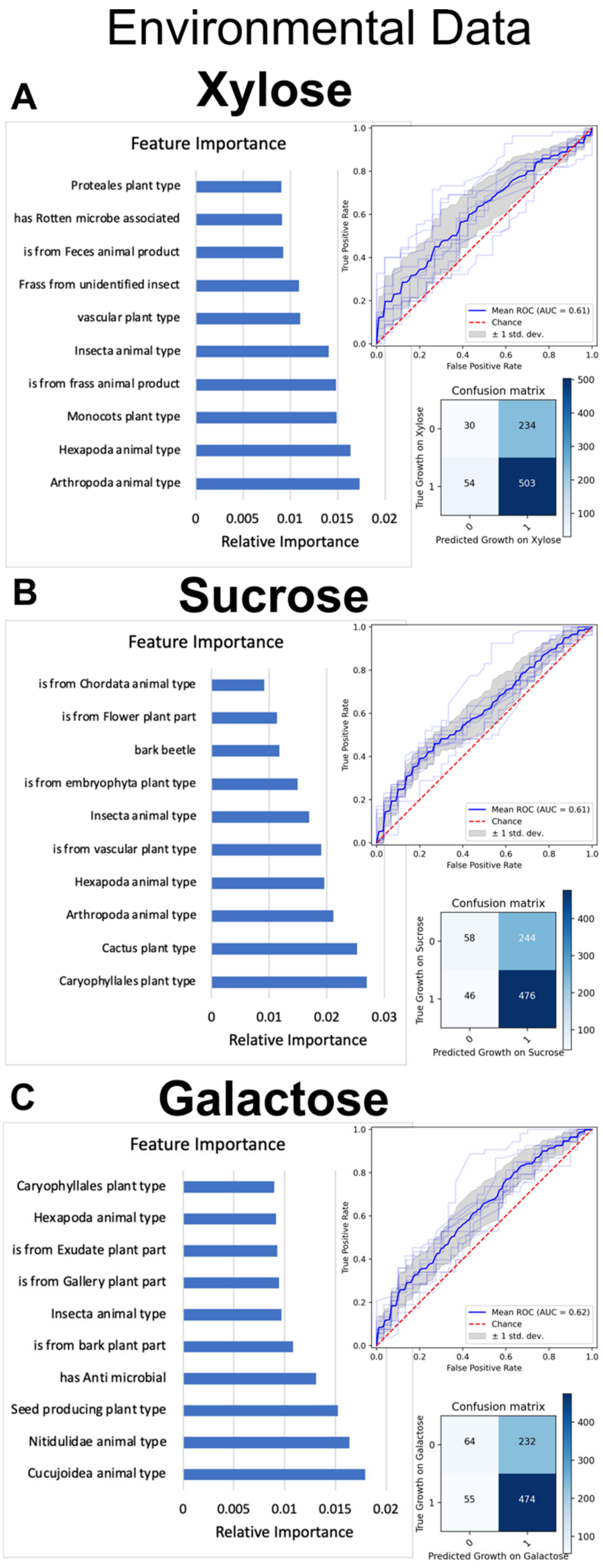
Environmental Data
A
Xylose
Feature Importance
Proteales plant type
has Rotten microbe associated
is from Feces animal product
Frass from unidentified insect
vascular plant type
Insecta animal type
is from frass animal product
Monocots plant type
Hexapoda animal type
Arthropoda animal type
0 0.005 0.01 0.015 0.02
Relative Importance
True Positive Rate
False Positive Rate
Mean ROC (AUC = 0.61)
Chance
± 1 std. dev.
Confusion matrix
30 234
54 503
True Growth on Xylose
Predicted Growth on Xylose
B
Sucrose
Feature Importance
is from Chordata animal type
is from Flower plant part
bark beetle
is from embryophyta plant type
Insecta animal type
is from vascular plant type
Hexapoda animal type
Arthropoda animal type
Cactus plant type
Caryophyllales plant type
0 0.01 0.02 0.03
Relative Importance
True Positive Rate
False Positive Rate
Mean ROC (AUC = 0.61)
Chance
± 1 std. dev.
Confusion matrix
58 244
46 476
True Growth on Sucrose
Predicted Growth on Sucrose
C
Galactose
Feature Importance
Caryophyllales plant type
Hexapoda animal type
is from Exudate plant part
is from Gallery plant part
Insecta animal type
is from bark plant part
has Anti microbial
Seed producing plant type
Nitidulidae animal type
Cucujoidea animal type
0 0.005 0.01 0.015 0.02
Relative Importance
True Positive Rate
False Positive Rate
Mean ROC (AUC = 0.62)
Chance
± 1 std. dev.
Confusion matrix
64 232
55 474
True Growth on Galactose
Predicted Growth on Galactose

**Supplemental Figure 1. Prediction accuracy of growth on different sugars was low when the random forest algorithm was trained on environmental data.** Prediction of growth on xylose (panel **A**), sucrose (panel **B**), and galactose (panel **C**) from environmental data. The right side of each panel shows the relative importance of different features (feature importance), which ranks the input features that are most useful for predicting growth on a given sugar. The top right graph of each panel is the Receiver Operating Characteristic (ROC) curve, which shows the true positive rate over false positive rate with changing classification thresholds. At the bottom right of each panel is the accuracy of classification in the form of a confusion matrix. Each confusion matrix shows strains predicted correctly to not grow on the sugar (true negatives, top left), strains predicted to grow on the sugar that do not (false positives, top right), strains correctly predicted to grow on the sugar (true positives, bottom right), and strains predicted to not grow on the sugar that do (false negatives, bottom left). Xgboost random forest was used to generate feature importance, and cross_val_predict() from sklearn.model_selection was used to generate confusion matrices. ROC curves were generated using the roc_curve function from sklearn.metrics.

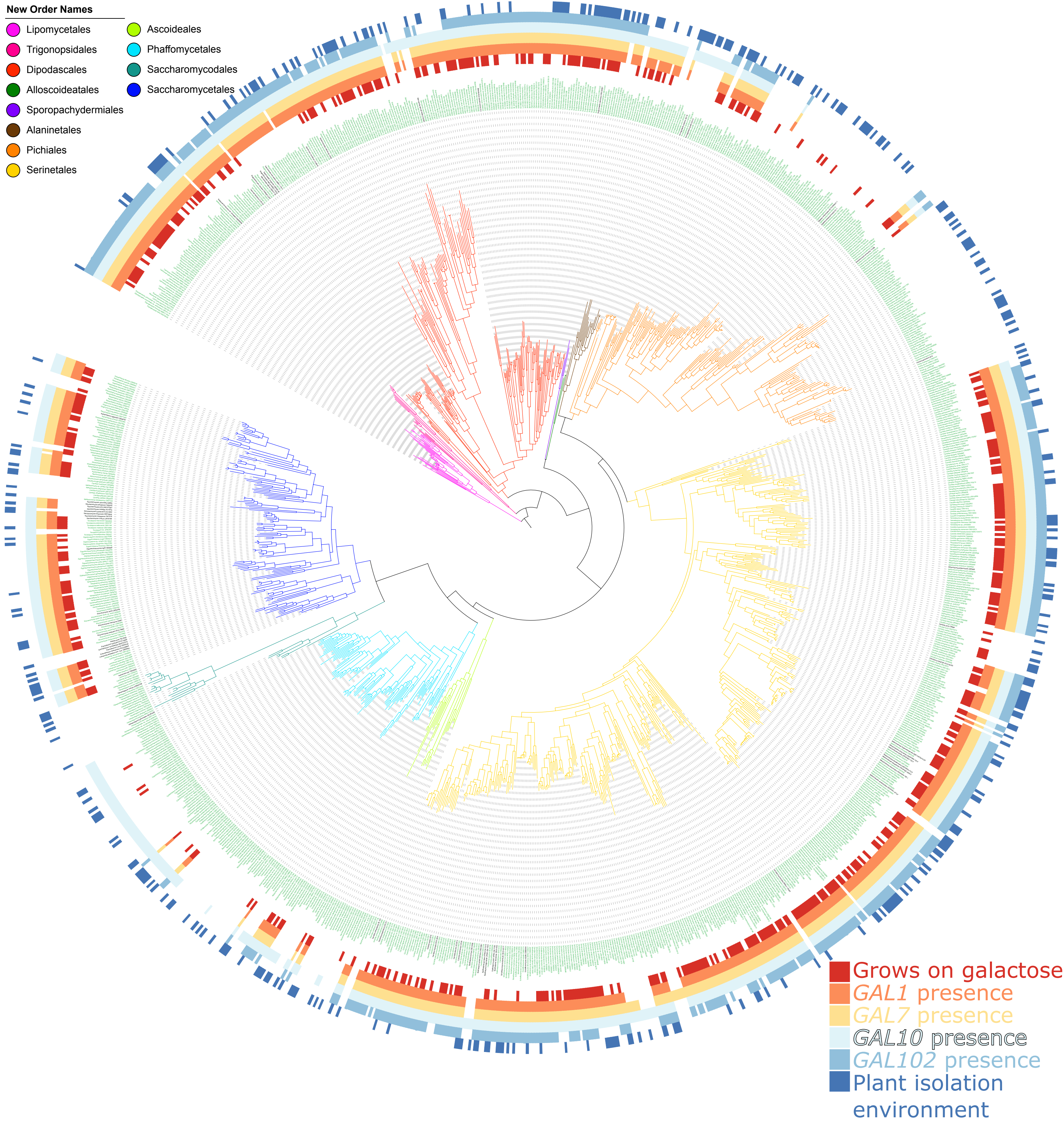
New Order Names
Lipomycetales
Trigonopsidales
Dipodascales
Alloscoideatales
Sporopachydermiales
Alaninetales
Pichiales
Serinetales
Ascoideales
Phaffomycetales
Saccharomycodales
Saccharomycetales
Grows on galactose
GAL1 presence
GAL7 presence
GAL10 presence
GAL102 presence
Plant isolation environment

**Supplemental Figure 2. Distribution of *GAL* genes and plant isolation environments across the Saccharomycotina phylogeny.** The ability of the different strains to grow on galactose, the presence of genes *GAL1*, *GAL7*, *GAL10*, and *GAL102*, and whether they were isolated from plant environments are plotted as circles (from innermost to outermost) around the Saccharomycotina phylogeny. The colors of the different branches of the Saccharomycotina phylogeny correspond to the 12 taxonomic orders(Groenewald et al. 2023).

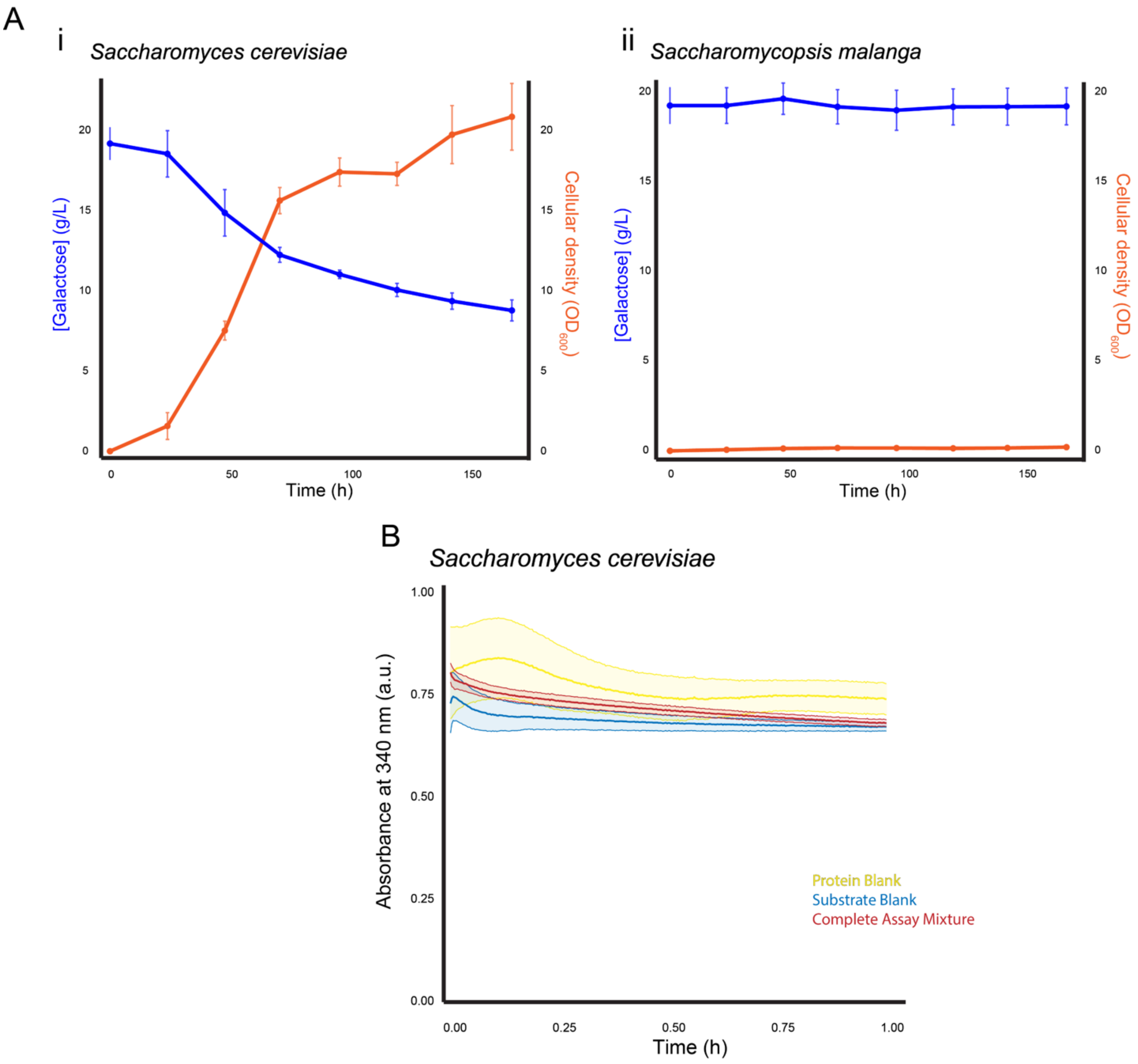


**Supplementary Figure 3. Positive and negative control data for experiments in Figure 7. A.** Average and standard deviation across three biological replicates of galactose concentrations in the medium (blue) and $OD_{600}$ growth measurements (orange) for the positive control species *S. cerevisiae* (i) and the negative control species *Saccharomycopsis malanga* (ii). **B.** Average and standard deviation across four biological replicates of NADPH absorbance at 340 nm over time for the negative control *S. cerevisiae* (red), the substrate blank for the negative control (blue), and protein blank for all species (yellow). The same protein blanks were used for all species included in the enzyme assay since each replicate of the enzyme assay included all four species on one 96-well plate, and the protein blank possessed reagents that were the same across all species (Tris-HCl, galactose, NADPH, and deionized water). Note that *S. cerevisiae* is a positive control for growth (Panel A) and a negative control for galactose reductase activity (Panel B).
